# Ecosystem stability relies on diversity difference between trophic levels

**DOI:** 10.1101/2024.08.23.609466

**Authors:** Yizhou Liu, Jiliang Hu, Jeff Gore

## Abstract

The stability of ecological communities has a profound impact on humans, ranging from individual health influenced by the microbiome to ecosystem services provided by fisheries. A long-standing goal of ecology is the elucidation of the interplay between biodiversity and ecosystem stability, with some ecologists warning of instability due to loss of species diversity while others arguing that greater diversity will instead lead to instability. Here, by considering a minimal two-level ecosystem with multiple predator and prey species, we show that stability does not depend on absolute diversity but rather on diversity differences between levels. We discovered that increasing diversity in either level first destabilizes but then stabilizes the community (i.e., a re-entrant stability transition). We therefore find that it is the diversity difference between levels that is the key to stability, with the least stable communities having similar diversities in different levels. An analytical stability criterion is derived, demonstrating quantitatively that the critical diversity difference is determined by the correlation between how one level affects another and how it is affected in turn. Our stability criterion also applies to consumer-resource models with other forms of interaction such as cross-feeding. Finally, we show that stability depends on diversity differences in ecosystems with three trophic levels. Our finding of a non-monotonic dependence of stability on diversity provides a natural explanation for the variety of diversity-stability relationships reported in the literature, and emphasizes the significance of level structure in predicting complex community behaviors.

## Introduction

Natural communities, from a microbiome to a forest, have a profound impact on humanity, influencing individual well-being and the sustainable development of societies^1,2^. Within these ecosystems, the interplay among millions of organisms from numerous species both within and between trophic levels^3,4^—such as fighting for territories^5^, predation^6–8^, competing for resources^9–11^, and cross-feeding^12–15^—creates a wide range of dynamic behaviors. These behaviors span a spectrum from global stability^16–18^ to rich, complex dynamics that include multi-stability^19–24^, periodic oscillations^6–8^, and even chaotic fluctuations^25–27^. Recognizing the interdependence of biodiversity and ecosystem stability, ecologists have spent decades striving to decode the mechanisms that enable complex ecosystems to achieve and sustain robust stability and functioning. Their works seek to illuminate how these myriad interactions contribute to the resilience of natural communities, which in turn impacts the stability of the environments we depend on.

Since Robert May’s pioneering work 50 years ago^28^, theoretical ecologists have used inter-species interaction strength and biodiversity to identify instabilities of large communities. May’s result predicts that communities will be destabilized by either strong interactions between species or a large number of species^28^. Subsequent research, employing generalized Lotka-Volterra (gLV) models, delved deeper into ecological networks without specifying mechanisms of interactions^29– 31^, examining how interaction patterns, such as the proportion of mutualistic versus competitive relationships^29,30^, impact the stability of ecosystems. One of the main takeaways was that increasing diversity is still expected to destabilize ecosystems, a robust theoretical expectation that is potentially at odds with empirical results that diversity can either stabilize or destabilize communities^32–40^. Recent investigations have revealed that higher-order interactions^41,42^, constraints on interactions^31,43^, or self-regulation^44,45^ can cause increased diversity to stabilize ecosystems. However, the primarily phenomenological nature of these models constrains our ability to correlate the mathematical parameters with tangible biological processes or environmental factors, limiting our predictions and understanding for real communities. We therefore look for understanding of ecosystem stability and its relationship to diversity through specifying mechanisms and structures of interactions.

In particular, predation or resource competition and the corresponding trophic level structures are important in real-world communities, whose impact to diversity-stability relationships is relatively poorly understood^3,46^. Predators may have only limited direct interactions but may instead compete primarily through competition for prey at a lower trophic level. More generally, a food web can include abiotic resources as a level in addition to trophic levels, where, for example, bacteria near the bottom of a food web may primarily compete for abiotic resources such as sugars. In considering the specific mechanisms like resource competition^47–57^, predation^58–61^, cross-feeding^12,47,52,55^, pH^27,62^, etc., the models incorporate key aspects of natural ecosystems but in many cases cannot be analyzed analytically. A variety of models have suggested that cross-feeding^50,52,55^, variation in yields^48,54,63,64^ (how efficient prey/resources are converted to predator/consumer biomass), and toxicity^47^ can destabilize communities, while some mechanisms like predator interference^59^ can stabilize communities. Stability criteria based on deterministic matrices successfully predicts the existence of instability due to various mechanisms^50,52,55^. However, we have not found a unified stability criterion based on simple community statistics, let alone understood how diversity affects stability.

Here, we focus on a minimal ecosystem model with two levels that could describe either predator-prey or consumer-resource interactions. In the context of predator-prey interactions these would typically be considered two trophic levels, whereas for consumer species growing on abiotic resources these would be the two lowest levels of a food web. We discovered a re-entrant stability transition in which increasing the species diversity in either level first destabilizes but then later stabilizes the community, which is absent in models without trophic levels. We then analytically derive a general stability criterion that quantitatively agrees with numerical results, revealing that stability prevails when the diversity difference between levels is substantial, irrespective of total diversity of the ecosystem. The critical diversity difference is determined by the correlation between how one level affects another and how it is affected in turn. We also find that mechanisms beyond predation or resource competition such as cross-feeding (production of resources by consumers) can destabilize the community, yet the stability criterion that we derived nonetheless correctly predicts the stability boundary. Beyond two-level ecosystems, numerical results further show that diversity difference remains the key determinant of stability in three-level ecosystems, suggesting that diversity differences between levels may play an important role in many ecosystems. Our finding of a non-monotonic dependence of stability on diversity provides a natural explanation for the variety of diversity-stability relationships reported in the literature. Our work emphasizes the significance of level structure in predicting community behaviors, and provides new insights into ecosystem stability.

## Results

To study the stability of an ecosystem with two levels we consider a simple model of predators (or consumers) in the higher level that grow based on consumption of the prey (or resources) in the lower level. We employ the dynamics for the predator abundance (*S*_*i*_) and prey abundance (*R*_*α*_) given by^47,49,52,56,59^:

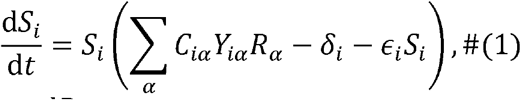

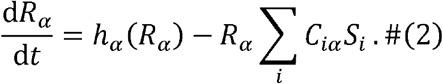

Here, *h*_*α*_(*R*_*α*_) refers to how prey grow, i.e., logistic growth *h*_*α*_(*R*_*α*_) = *g*_*α*_ *R*_*α*_(*K*_*α*_ − *R*_*α*_), or how resources are supplied to the system, i.e., chemostat supply *h*_*α*_(*R*_*α*_) = *l*_*α*_ (*κ*_*α*_ − *R*_*α*_). Our conclusions are valid for either functional form (See Methods) and therefore we do not emphasize the function *h*_*α*_ in the text. Numerical results in the main text are obtained from logistic growth while we present the counterparts (see Fig. S1) for chemostat in supplementary information (SI). The above equations describe only predation or resource competition (Fig. 1a), but we can generalize it to include cross-feeding and resource-resource regulation under the consumer-resource picture (see Methods). *C*_*iα*_ describes the per capita consumption rate for predator (species) *i* on unit prey (species) *α*. Prey consumed will not be fully converted to biomass of predator, so *Y*_*iα*_ quantifies the associated yield (negative *Y*_*iα*_ could then capture toxicity). We aim to study the stability of the ecosystem at a fixed point, and the linear consumption and growth can then be thought of as local linear approximations of some complicated consumption and growth functions. Without loss of generality, we assume the fixed point of interest has *N* predators and *M* prey and focus on their dynamics. Finally, predators are set to have mortality *δ*_*i*_ and autoregulation *ϵ*_*i*_. In the canonical consumer-resource models, there is no autoregulation for consumers (*ϵ*_*i*_ = 0)^49,50,52,55^, while in predator-prey models, predator autoregulation is common^59–61^. We include autoregulation for generality, which also overcomes the competitive exclusion principle^65–67^ (i.e., the number of predators *N* that can coexist being limited by the number of prey *M*). We initially assume predator autoregulation is much smaller than other interactions for analyticity but later discuss how predator autoregulation will shift the analytic results.

**Fig. 1.**
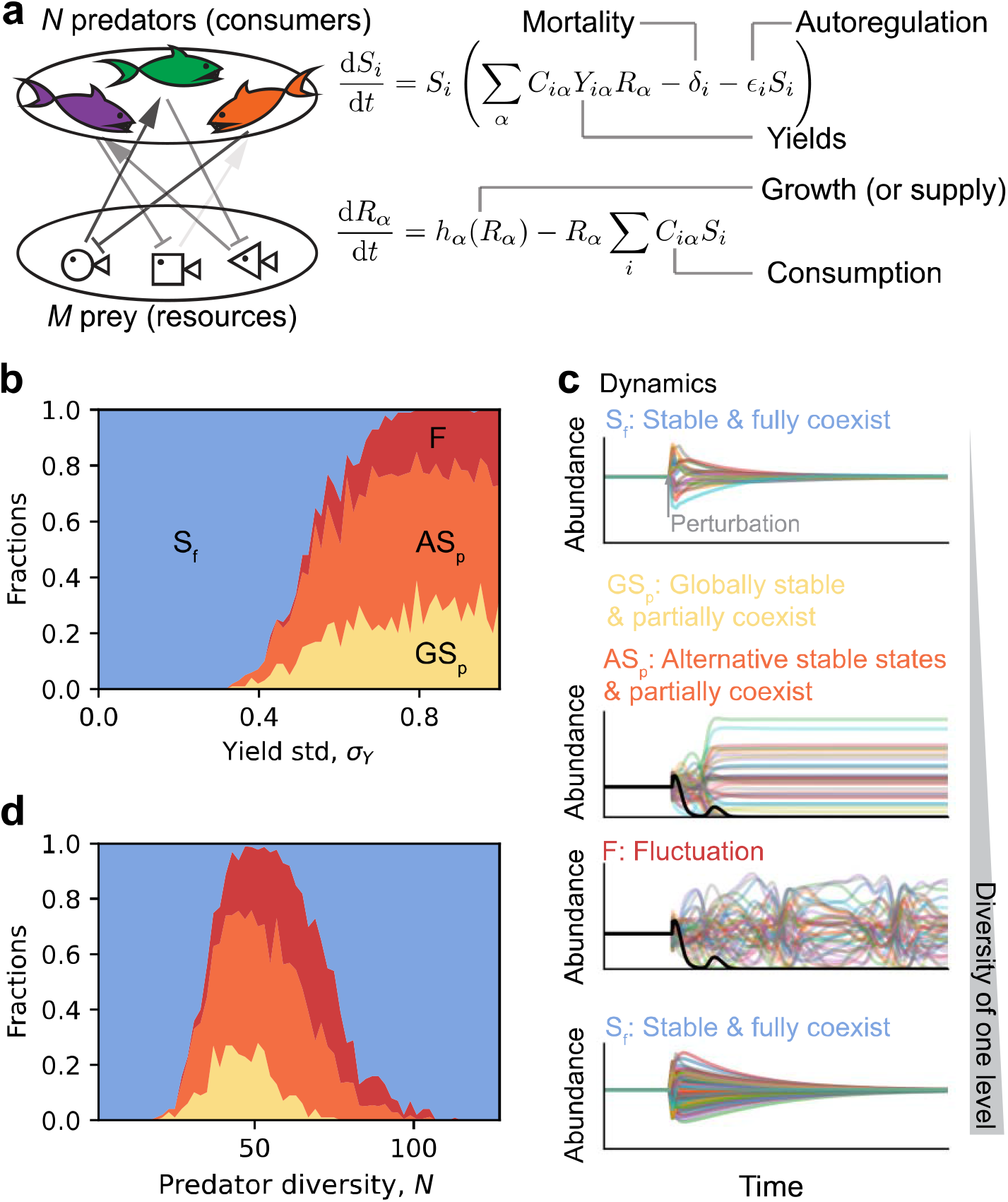
Two-level ecosystems that could describe either predator-prey or consumer-resource dynamics are first destabilized but then stabilized by increasing predator (or prey) diversity at the fixed point. (a) A minimum model with *N* predator species and *M* prey species. Consumption (*C*_*α*_) depletes prey while promotes predator growth. Prey consumed transfer to predator biomass increasement subject to yields (*Y*_*iα*_). Prey growth (or resource supply) is governed by *h*_*α*_ function. Predator death is encoded in mortality rates (*δ*_*i*_). And predators may have autoregulation (*ϵ*_*i*_) limiting growth. (b) Only changing the variance of yields, we observe one transition to instability with multiple possible behaviors after the loss of stability of the original fixed point (S_f_): the community can be globally stable but with extinction (GS_p_), or it can go to alternative stable states with partial coexistence, or it can display persistent fluctuations (F). We fixed prey diversity *M* = 64 and predator diversity *N* = 32 (details in SI). (c) Time series examples for different dynamical behaviors. We plot the abundances normalized by their original fixed-point values. At first, the communities are at the given fixed point. Later, communities are perturbed (starting from another randomly sampled initial condition). If the time series come back to the given fixed-point values, the community is S_f_. GS_p_ and AS_p_ share one illustrative time series as they all end up being stable but having extinctions (highlighted by the dark curve). (d) Only varying predator diversity, *N*, the fraction of S_f_ first decreases but then increases with *N* (re-entrant stability transition). The color encoding for different dynamical behaviors is the same as (b). We fixed prey diversity *M* = 48 and varied *N* from 1 to 128, and the fractions of different dynamical behaviors in the figure were averaged over communities with the same *N* (details in SI).

In a large community we expect only the statistical properties of parameters to determine the stability. Since the growth rate of a predator cannot go to infinity when there are more available prey, individual consumption rates *C*_*iα*_ should scale as 1/*M*. We therefore sample *C*_*iα*_ with mean and standard deviation being *µ*_*C*_ /*M* and *σ*_*C*_ /*M*, respectively (the actual distribution does not influence the results, but main text figures are obtained with a uniform distribution). The yields are determined by biochemical processes and should not scale with community size. We sample yields, *Y*_*iα*_, from a Gaussian with mean, *µ*_*Y*_, and standard deviation, *σ*_*Y*_. If not specified, we will use *µ* and *σ* to denote means and standard deviations of other parameters, with proper subscript showing which parameter we are referring to.

Although the original MacArthur (consumer-resource) model only displays global stability^56^, allowing yields to vary depending on both predators and prey can lead to the loss of stability^48,54,63,64^. Predators (consumers) may digest prey (resources) differently and prey (resources) can contain nutrients differently, which leads to yield variation observed^48^. One way to understand why variation in yields destabilizes the community is that it leads to “niche encroachment”, whereby predators consume prey important for the growth of other predators^54^. Indeed, sampling yields as i.i.d. and increasing the standard deviation, *σ*_*Y*_, causes a loss of stability and a variety of community outcomes (Fig. 1b, numerical details in SI). Some communities remain globally stable but with some extinction, whereas other communities display sustained fluctuations in abundance (limit cycles and chaos) or alternative stable states with different community compositions (Fig. 1c). Based on the idea of niche encroachment, we would expect that, holding the variation in yields constant, increasing the number of predators will make the niches more crowded and therefore lead to instability.

To test this expectation, we studied how diversity of the top level (number of predator species, *N*) affects community stability. We fixed other statistical parameters (where *σ*_*Y*_ ≠ 0) and prey diversity (number of prey species, *M*) when sampling communities but varied predator diversity, *N*, and then simulated the dynamics of each sampled community (details in SI). With a small number of predators, all communities are globally stable at the original fixed point where all N predators survive (Fig. 1d). As expected, increasing predator diversity can lead to the loss of stability of the original fixed point and the full range of rich dynamical behaviors discussed previously. Although increasing the number of predators in the top level led to a loss in stability, we were surprised to find that further increases in predator diversity led the community to regain stability (Fig. 1b). The observation that stability depends non-monotonically on predator diversity contradicts the idea that more predator species leads to more niche encroachment and only destabilize communities. We therefore observed a counterintuitive re-entrant stability transition with respect to predator diversity, i.e., increasing predator diversity first destabilizes but then stabilizes the communities.

Having characterized community outcomes after varying the diversity of the top level (number of predator species), we next sought to characterize community outcomes when varying the diversity of the lower level (number of prey species). We therefore sampled communities only varying prey diversity, □, and again found a re-entrant stability transition in which communities lost but then regained stability (SI, Fig. S2). Our observation that stability depends non-monotonically on prey diversity also contradicts the idea that more prey species provide more niches and only stabilize communities.

To gain insight into the re-entrant stability transition that we observed, we next varied predator diversity *N* and prey diversity *M* while holding other statistics unchanged. We observed that the fraction of unstable communities depends differently on predator diversity *N* for different prey diversity *M* (Fig. 2, a and b). Nevertheless, instability depends on *N* and *M* in a similar way, with the most unstable cases being when the diversities of the two levels are similar (*N* ≈ *M*). Surprisingly, after we normalize the predator diversity by prey diversity, we found a universal pattern that increasing predator-prey ratio, *N*/*M*, first destabilizes but then stabilizes communities, giving two stability-instability boundaries (Fig. 2c). The stability of our two-level community therefore depends fundamentally on the difference in diversities of the two levels, with the community being least stable when the diversities in the two levels are similar.

**Fig. 2.**
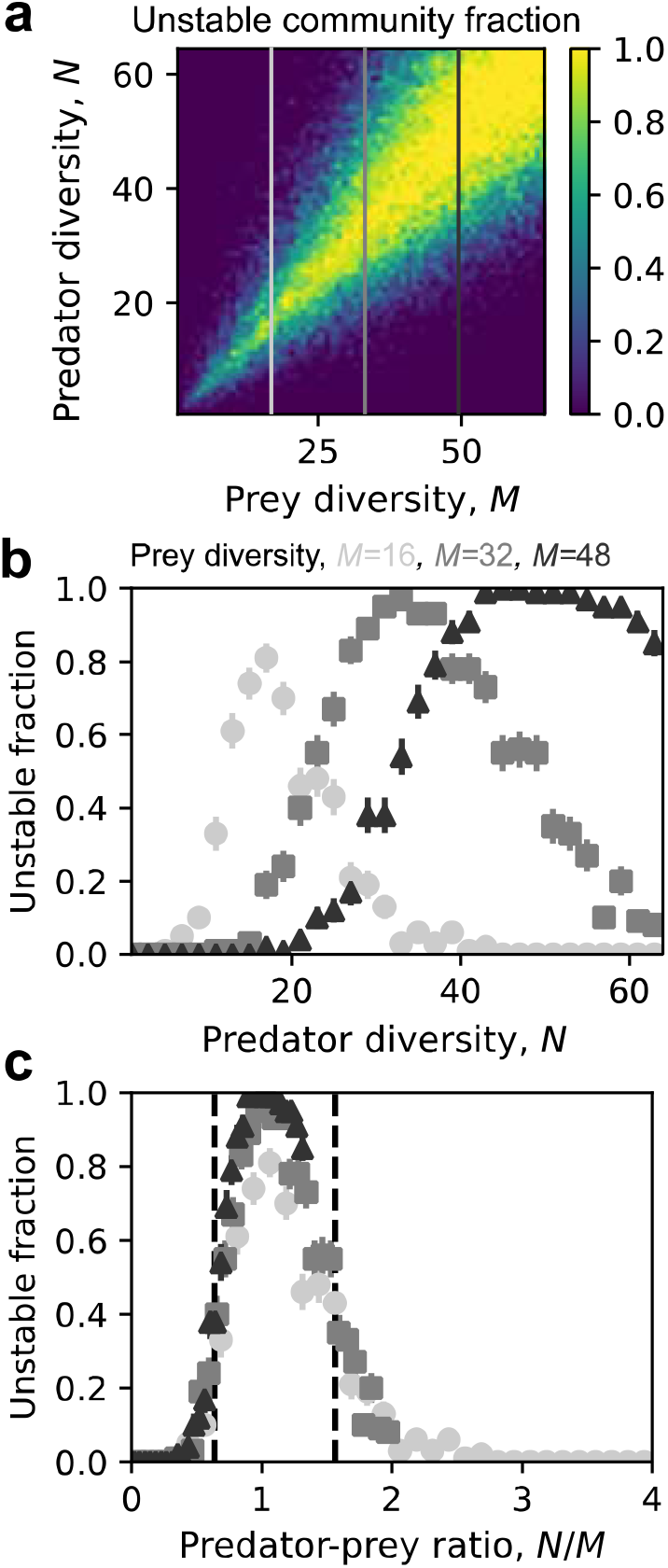
Stability depends on diversity difference between trophic levels, with a greater difference leading to stability. (a) For communities with different number of prey, stability depends differently on predator diversity, but the fraction of unstable communities seems to have a peak near *N* = *M*, indicating the universal importance of *N*/*M*. The color gradient represents the fraction of unstable communities, with higher values indicating a greater fraction of instability. The vertical lines indicate different levels of prey diversity (*M* = 16, 32, 48). (b) Fraction of unstable communities are plotted in more detail for selected prey diversity (*M* = 16, 32, 48), which shows different dependence on predator diversity. (c) There is indeed a universal trend of stability for different communities if we vary the ratio of number of predators to that of prey, *N*/*M*: communities will first be destabilized but then stabilized (re-entrant stability transition). (b) and (c) used the same set of data. Each data point in the figure was averaged over 100 sampled communities. Error bars represent standard error of mean (SEM).

We next sought to explain the surprising re-entrant stability transition that we observed with respect to predator-prey ratio. We followed the standard stability analysis to study the Jacobian at the fixed point (Fig. 3a). If we combine predator and prey abundances into one vector, the Jacobian becomes a block matrix, where in our convention the upper right part, Λ (an *N* × *M* matrix), encodes how prey affect predators, the lower left part, *V*^*T*^ (an *M* × *N* matrix), encodes how predators modify prey, and the lower right part is an *M* × *N* diagonal matrix encoding effective autoregulation of prey (see Methods). In the regime *N* < *M*, with the proper approach of calculating block matrix determinant, we can have *M* − *N* stable eigenvalues directly from prey autoregulation, and solve the rest of 2*N* eigenvalues from eigenvalues of Λ*V*^*T*^ via quadratic equations (see Methods). Since Λ*V*^*T*^ is an *N* × *N* matrix passing the effects from predator-changed prey back to predators (Fig. 3b), it is naturally understood as (reduced) inter-predator interactions. In large complex systems, we can show that whether the Jacobian can have unstable eigenvalues depends on whether Λ*V*^*T*^ has unstable eigenvalues (Methods). In other words, stability of the large community is determined by reduced inter-predator interactions when *N* < *M*. By symmetry, one can imagine that in the regime *N* > *M*, stability is determined by reduced inter-prey interactions, *V*^*T*^Λ (Fig. 3b), which we can prove (see Methods). Symmetry or duality therefore suggests that there will be two instability transitions, with the transition at *N* < *M* determined by inter-predator interactions (Λ*V*^*T*^) and the transition at *N* > *M* determined by inter-prey interactions (*V*^*T*^Λ).

**Fig. 3.**
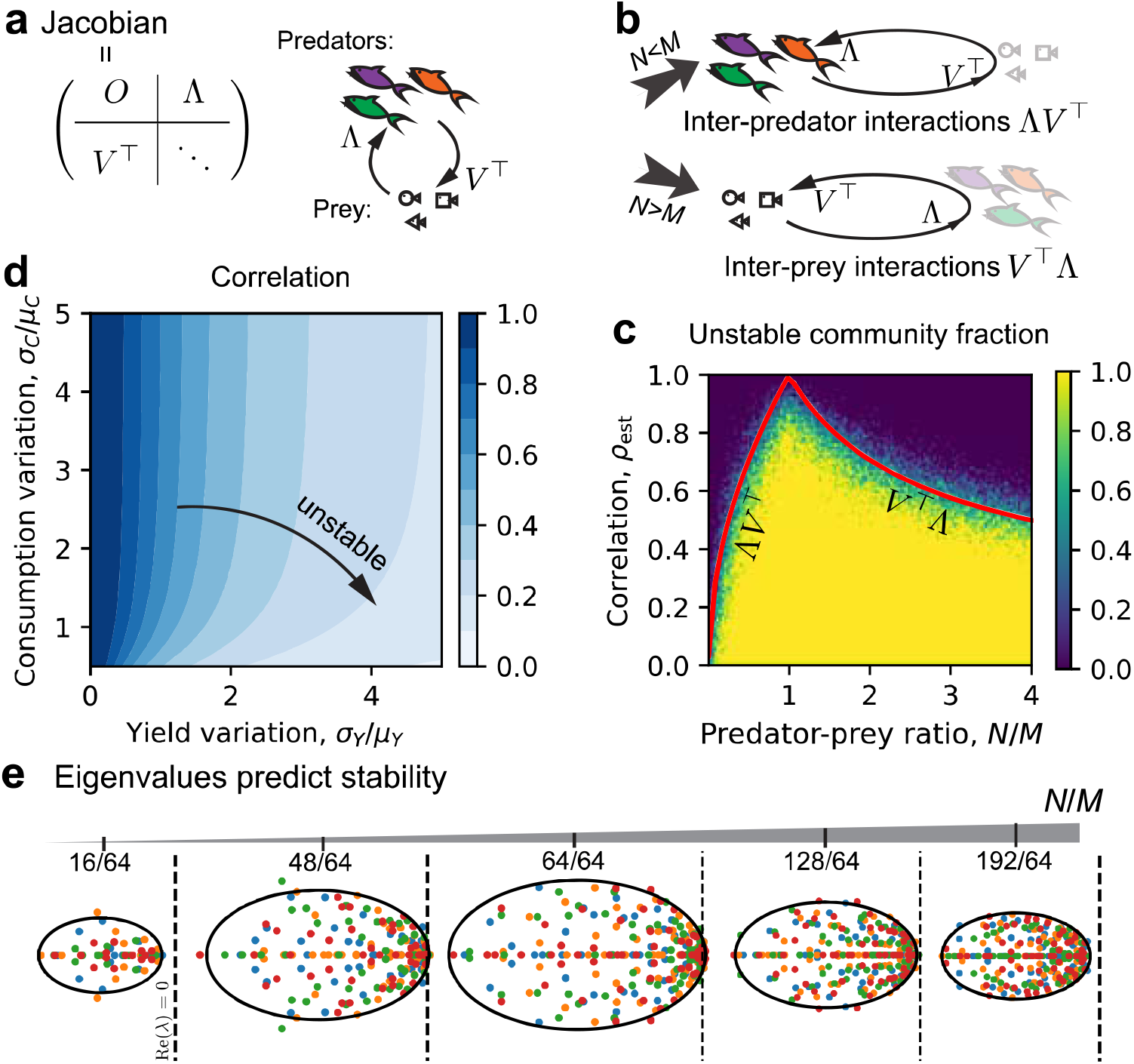
Stability relies on the predator-prey ratio and the correlation between how predators affect prey and how prey in turn influence predator. (a) The local Jacobian at the fixed point with negligible predator autoregulation is a block matrix mainly consist of how prey affect predators (the matrix Λ), how predators affect prey (the matrix *V*^*T*^), and effective prey autoregulation (lower right diagonal matrix). (b) When there are more prey than predators (*N* < *M*), stability depends on the matrix Λ*V*^*T*^ (reduced inter-predator interactions) and effective autoregulation, and will be determined only by the stability of Λ*V*^*T*^ in large communities. When there are more predators than prey (*N* > *M*), similarly, stability for large communities will be determined by that of the matrix *V*^*T*^ Λ (reduced inter-prey interactions). (c) In different regimes (*N* < *M* and > *M*), the stability boundaries are derived (red lines), depending on predator-prey ratio and the correlation between Λ and -*V* elements (estimated here from mechanistic parameters), agreeing well with numerical results. The heatmap is obtained from numerically sampled communities with *M*= 32 and *N* varying from 1 to 128. There are 128 × 100 pixels, and each pixel has 10 communities fell in, from which the unstable community fraction can be calculated. (d) The estimated correlation (Eq. (6), which is exact when yields and consumption are i.i.d., respectively) depends on the variation in yields, *σ*_*Y*_ /*μ*_*Y*_, and the variation in consumption, *σ*_*C*_ /*μ*_*C*_. (e) The eigenvalues of Λ*V*^*T*^ (or *V*^*T*^Λ when *N*/*M* > 1) are in an ellipse in the complex plane, and stability depends on if the ellipse goes beyond the dashed line (The dashed lines mark where the real part is zero; note the real axis is horizontal and imaginary axis is vertical by convention).

After simplifying the stability analysis to studying inter-predator or inter-prey interactions, we then could derive an analytic and complete stability criterion. For large communities, Λ*V*^*T*^ (or *V*^*T*^Λ) can be viewed as a large random matrix and we can apply the non-Hermitian Marchenko-Pastur law^54,68,69^ (see Methods). The eigenvalues of Λ*V*^*T*^ are then distributed in an ellipse in the complex plane (up to a non-essential normalization not changing signs of eigenvalues). The ellipse has a center on the real axis at − *ρ*(1 + *N*/*M*), whose major axis is also on the real axis and has length 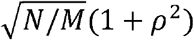, where − *ρ* is the correlation between Λ_*iα*_ and *V*_*iα*_ (Fig. 3e). Stability requires that all eigenvalues have negative real part, leading to the stability criterion when *N* < *M* given by

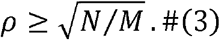

For *V*^*T*^Λ, we only need to exchange *N* and *M* in the derivations related to Λ*V*^*T*^ and can directly obtain the criterion for *N* > *M* given by

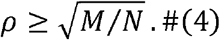

A concise form combining the two cases then given the complete stability criterion as

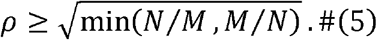

Random matrix analysis therefore allows for a simple stability criterion that depends upon how the ratio of diversities in the two levels compared to the correlation in interactions between the levels.

We next explored the meaning of the inter-level correlation *ρ*, and how it is affected by mechanistic parameters connecting the predators and prey. As a general statement, the correlation *ρ* measures the alignment between how predators affect prey and how prey in turn influence predators. This statistical parameter quantifies the degree of reciprocity or feedback between these two levels. For example, if a predator species kills certain prey at high rates but cannot grow well on that prey, the correlation *ρ* is reduced. On the other hand, if all yields are uniform and positive then the correlation is maximized as 1.We estimated the correlation when yields have variation as (see Methods for derivation)

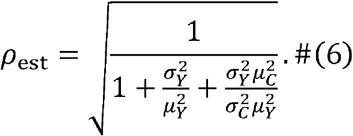

With this estimated correlation, we have a stability criterion that purely depends on mechanistic parameter properties and diversities, which is well supported by numerical experiments (Fig. 3c, where the red line is theoretical stability boundary and the heatmap is numerical results on stability). For a community with given sizes (*N* and *M*), greater variation in yields, quantified by *σ*_*Y*_ / *μ*_*Y*_, or smaller variation in consumption, *σ*_*C*_ / *μ*_*C*_, can reduce the correlation between predators affecting prey and prey impacting predators, tending to destabilize the community (Fig. 3d). A greater yield variation makes predators more likely to encroach upon growth promoting prey (niches) of others and thus tend to result in instability^54^. Similarly, a small consumption variation means predators are competing for a very similar set of prey, leading to strong interactions and subsequently, instability. We have therefore rationalized re-entrant stability transitions via the symmetric roles of inter-predator and inter-prey interactions, identified a critical diversity difference for instability controlled by correlation, and interpreted this correlation as the alignment between inter-level interactions or how well species focus on their own niches.

Beyond cases we could analyze analytically, we further explored the generality of our findings through numerical tests. Natural ecosystems may have more than two levels in the food web, where our results cannot be directly applied. To gain some insight into more complex ecosystems, we studied stability of a three-level community containing *M* prey species, *N*_1_ predator species on those prey, and *N*_2_ apex predators that consume the middle predators (Fig. 4a). We assumed that there are only direct interactions between neighboring levels and the correlations of inter-level interactions are the same (see details in SI). The stability is found to depend on both diversity ratios, *M*/*N*_1_ and *N*_2_/*N*_1_, with the most unstable cases having *M* + *N*_2_ ≈ *N*_1_ (Fig. 4b). This finding inspired us to plot the stability as a function of diversity difference (*M* + *N*_2_)/*N*_1_, which then show a unified re-entrant transition (Fig. 4c). These results suggest that diversity difference between trophic levels may therefore play a special role in determining stability even in multi-level ecosystems.

**Fig. 4.**
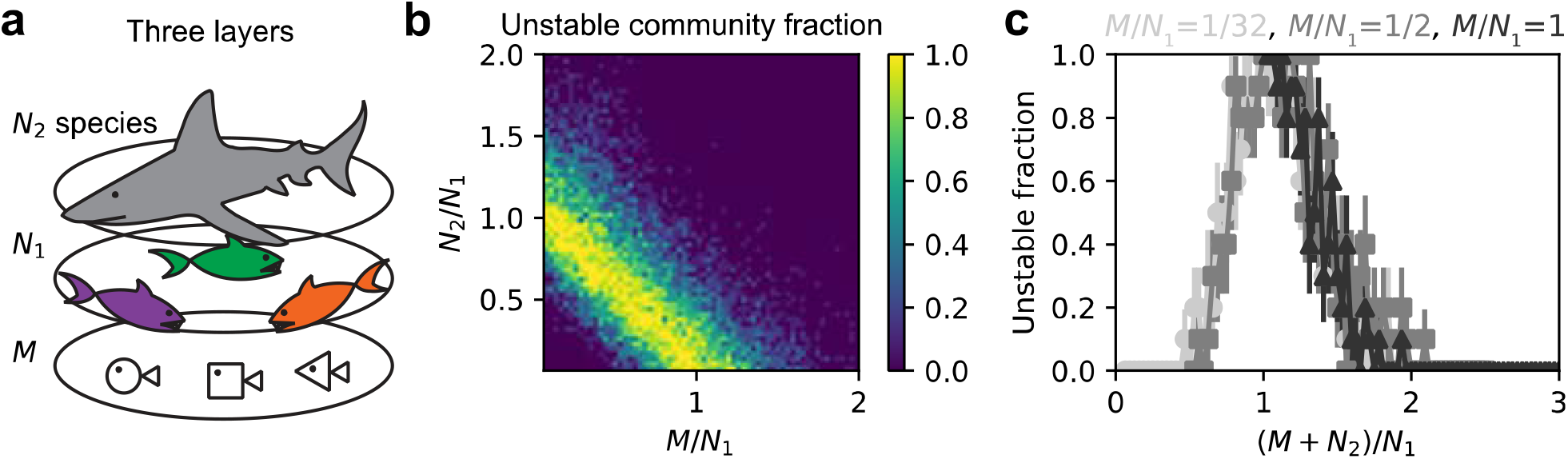
Numerical evidence suggests that stability also depends on diversity differences in communities with three levels. (a) Illustration of a three-level community has *M* species in the bottom level, *N*_l_ species in the middle level, and *N*_2_ species in the top level. (b) For one instance, we found stability depends non-trivially on both ratios of diversities (of nearby levels), *M*/*N*_l_ and *N*_2_/*N*_l_, with the most unstable cases having *M* + *N*_2_ ≈ *N*_l_. We kept *N*_l_ = 32, while changed *M* and *N*_2_ from 1 to 64, respectively. (c) The diversity difference quantified by (*M* + *N*_2_)/ *N*_l_ seems to determine stability transitions, as different re-entrant transitions collapse if we fraction of unstable communities with respect to (*M* + *N*_2_)/ *N*_l_. Data in (c) are from (b). Error bars represent SEM. We assumed for the instance shown that there is no direct interaction between top and bottom levels, inter-level correlations are all *ρ* = 0.8, and the middle level has negligible autoregulation as in conventional consumer-resource models. The multi-level communities need some levels to have sufficiently weak autoregulation to be unstable, and the instability seems to depend on difference between diversity of levels having negligible autoregulation and that of other levels (See more discussions and examples in SI).

## Discussion

We have studied a two-level ecosystem (predator vs. prey or consumer vs. resource) where interactions between species are due to predation or competition for food. The level structure leads to the qualitatively new phenomenon of a re-entrant stability transition: increasing diversity of either level first destabilizes but then stabilizes the communities. This observation of a return to an original phase (in this case stability) as a function of a control parameter is similar to re-entrant phase transitions observed in liquid crystals^70^, RNA-polycation mixtures^71^, and polypeptides^72^ but with very different mechanisms. However, the re-entrant phenomenon of “double descent”^73,74^ in machine learning, where more training data first leads to greater errors but then additional data leads to smaller errors, has a similar mathematical origin via random matrix theory, namely the Marchenko-Pastur law^75,76^. These re-entrant behaviors are counterintuitive as we do not expect a system to return to its original state when tuning one parameter monotonically, and more work is required to clarify the similarities and differences of the various types of re-entrant transitions.

In our communities, predator species in the higher level do not interact directly with each other, but instead all interactions are mediated by competition for the prey (or resources) at the lower level. When there are more prey than predators (*M* > *N*), it is the inter-predator interactions (Λ*V*^*T*^) that determine stability. Increasing the number of predators then destabilizes the community, consistent with previous results obtained in fully-connected Lotka-Volterra type models^28,29^. In fact, by heuristically comparing our stability criterion to May’s^28^, we can obtain an explicit expression for the interaction strength (derived in Methods: greater yield variation or fewer prey leads to stronger interaction strength). However, when we have more predators than prey (*N* > *M*), it is the inter-prey interaction (*V*^*T*^Λ) that determines stability, meaning that increasing predator diversity stabilizes the communities as it weakens the inter-prey interactions. The re-entrant stability transition is therefore due to this symmetric behavior between predators and prey that emerges in a two-level ecosystem.

The symmetry argument that when *N* > *M* we need to shift to consider inter-prey interactions is one straightforward way to understand the existence of re-entrant transitions yet does not present the complete picture. The matrices Λ*V*^*T*^ and *V*^*T*^Λ share the same non-zero eigenvalues, meaning that we can predict stability correctly with inter-predator interactions Λ*V*^*T*^ alone. The re-entrant stability transition may therefore arise from inter-predator or inter-prey interactions alone rather than via the transition from inter-predator interactions to inter-predator ones. A puzzle then emerges as the inter-predator interaction Λ*V*^*T*^ can always predict stability but the intuition for the inter-predator interaction, i.e., the niche encroachment, only works for communities with fewer predators than prey. Specifically, why does the intuition that adding predators leads to strong niche encroachment and therefore instabilities work when *N* < *M* but fail when *N* > *M*? Our naïve interpretation is that when *N* > *M*, inter-predator interactions are highly correlated (linearly dependent) as predators are competing for fewer prey, which not only leads to zero eigenvalues but also a stabilization effect. Further work is necessary to clarify the ecological intuition behind this stabilization effect that leads to the re-entrant stability transition.

The re-entrant stability transition can also be observed with a single-level modeling framework in which the two-level structure is contained within the effective interactions. For example, a gLV model containing only the predators but with the interaction matrix given by Λ*V*^*T*^ will also display the re-entrant stability transition with respect to predator diversity (where the number of prey species *M* is encoded in Λ*V*^*T*^). The two-level structure of the ecosystem that leads to the re-entrant stability transition is in this case encoded in the structure of the interaction matrix of the single-level model of predators.

In our model, we included predator autoregulation whereby increasing abundance of a predator species leads to inhibition of itself. This autoregulation could capture a variety of effects such as competition for territory^5^ or intra-species aggression^77^. If predator autoregulation is strictly zero, the number of predators is always smaller than that of prey (*N* < *M*) in the absence of fine-tuned metabolic trade-offs^67^ and we therefore cannot observe any re-entrant stability transition. However, stability still depends on diversity difference between levels (Eq. (3)). We assumed the predator autoregulation is much weaker than other mechanisms and therefore could derive a simple theoretical result for stability (Eq. (5)). If predator autoregulation is non-negligible it stabilizes communities, but we can still observe a re-entrant stability transition (SI, Fig. S4). Adding strong predator autoregulation therefore stabilizes communities but stability still depends on diversity difference between levels of a food web.

An important question is the robustness of the re-entrant stability transition and associated analysis that we have explained in a simple two-level community. As already shown, the argument that stability depends on diversity differences rather than total diversity still holds for ecosystems with more than two levels, yet the diversity difference can have distinct forms compared to the simple two-level case. Another direction to test robustness is to include more interactions besides predation or resource competition. It is known that cross-feeding under the consumer-resource picture plays an essential role in determining community structure^12–15^ and has been show to lead to instabilities^50,52,55^. Our analysis can include cross-feeding, and we found the stability criterion, Eq. (5), remains the same while the correlation *ρ* will be decreased: if one consumer is able to produce metabolites as resources which does not affect its growth rate much, the reciprocity between the two levels is reduced (see Methods and Fig. S3-S4). Cross-feeding in our framework therefore destabilizes communities quantitatively as predicted by its effect on *ρ*. In addition, we found that higher-order dependence of predator growth on prey abundances also destabilizes the communities in a similar manner (Methods and Fig. S3-S4). Our analytic analysis can therefore be generalized to capture the re-entrant stability transition that occurs under a range of interactions.

Other modifications to the model may alter the re-entrant stability transition in a more profound way. Here we focused on large ecosystems, where we found that stability is determined by inter-predator or inter-prey interactions. However, in small communities, stability can be lost due to too weak effective autoregulation of prey (SI, Fig. S5). An example is the original Lotka-Volterra predator-prey model^78,79^, where there is only one predator species and one prey species, and the inter-predator interaction is always stable (Λ*V*^*T*^ is a negative scalar number) but the fixed point is not stable. We studied communities lacking interactions within the same level, and the effects of including such intra-level interactions are not clear. Similarly, our preliminary results on three-level communities were obtained from particular examples, and generalizing to include more levels or predation across multiple levels may alter the dynamics (SI, Fig. S6). Future studies are therefore necessary to explore the generality and consequences of the re-entrant stability transition discovered here.

It has been a long-standing intuition that level structure in food webs may alter stability-diversity relationships^40^. Greater diversity can enhance stability by providing alternative pathways for energy flow in the food web, thus buffering against the loss of species. However, more species can also lead to instability due to strong interactions. In fact, increasing the number of prey species was observed to stabilize communities in some cases^33–35^ while to destabilize communities in other cases^36^. Without our theory suggesting that it is the ratio of diversities in different levels that is the key to stability, studies do not always quantify diversities in different food web levels systematically and record dynamical behaviors at the same time. Our theory then provides a possible explanation for the observations of both positive and negative stability-diversity relationships^32–40^. The theory may also provide insight into biodiversity patterns across trophic levels distinct from arguments like energy and mass flow^80^. The “paradox of the plankton”^81,82^ can be rationalized as stability under more consumers (predators) than resources (prey), although the number of resources is often unclear due to cross-feeding. Future work will therefore be required to determine whether the re-entrant stability transition predicted here is a significant force structuring natural communities.

## Acknowledgements

We thank all the Gore Laboratory members for their valuable suggestions.

## Author contributions

Y. L. conceptualized the study, conducted analysis, and wrote the manuscript; J. H. conceptualized the study and revised the manuscript; J. G. conceptualized the study and wrote the manuscript.

## Methods

We study the dynamics of predator abundances denoted by *S*_*i*_ and prey abundances denoted by *R*_*α*_. The time evolution is governed by

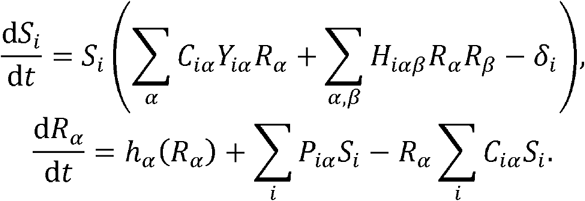

Say there is an equilibrium of the system with *N* predators and *M* prey given by *S*^*^ and *R*^*^ (two column vectors). Besides parameters introduced before, we here add high-order prey regulation for predator growth (*H*_*iαβ*_) and cross-feeding (allowing consumers to produce resource: *P*_*α*_) in terms of consumer-resource picture. There are all kinds of high order regulations that one prey may help or inhibit the consumption of another prey or itself. For simplicity, we choose *H*_*iαβ*_ to be mean zero (no community level trend for regulations to be beneficial or harmful). And we assume the *H*_*iαβ*_ to have standard deviation *σ*_*H*_/*M*^3/2^ such that if *σ*_*H*_ is constant, the total effect of regulation summed over all prey/resources has the same order of magnetite as the original growth rate (correction due to high-order interactions is not overwhelming). The production rates of resource or metabolites, *P*_*iα*_, are assumed to have mean *μ*_*P*_/*M* and standard deviation *σ*_*P*_/*M*, which follows similar logic to *C*_*iα*_: total production of different resources of one consumer individual is of constant order. We ignore invasion of other species not at the fixed point. The local Jacobian can be obtained by standard linearization at the fixed point:

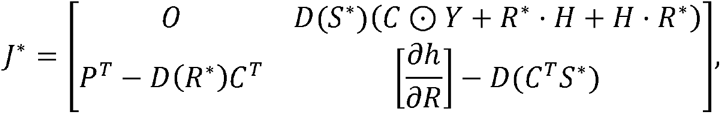

which is written as a block matrix if we concatenate predator vector and prey vector as [*S*; *R*]and consider its change around the fixed point. Here, we use *D*(*S*)^*^for the diagonal matrix with vector *S*^*\**^ on the diagonal, ⊙ to denote the Hadamard product, *R*^*\**^ · *H* to represent matrix 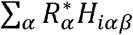, *H* · *R*^*\**^ for 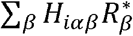 and 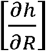 for the matrix with elements 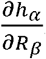.

For different supply/growth functions, the lower right block (effective prey autoregulation) is given as follows. For chemostat supply, *h*_*α*_(*R*_*α*_) = *l*_*α*_(*κ*_*α*_ − *R*_*α*_) and therefore

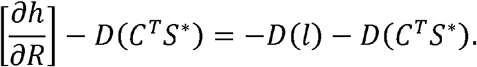

And for logistic growth, *h*_*α*_(*R*_*α*_) = *g*_*α*_ *R*_*α*_(*K*_*α*_ − *R*_*α*_), which gives

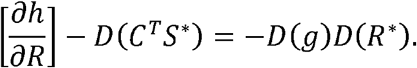

Note that in calculation, we have used the fixed-point property, 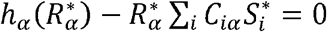. Regardless of different supply/growth functions, the lower right block is diagonal where all elements are negative (thus it represent autoregulation). Besides, each element in the diagonal is of the order of constant (not scaling with system sizes), which is important for later reasoning.

The stability of this Jacobian highly depends on the upper right block,

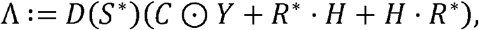

and the lower left one,

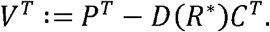

We consider an approximation where the lower right part, i.e., the effective autoregulation, is proportional to identity matrix, and rewrite the Jacobian as

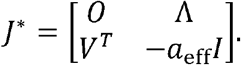

Since the lower right part should be negative, we restrict *a*_eff_ to be a positive number. Let *λ* ^*J*^ be the eigenvalue of the Jacobian, which can be solved from

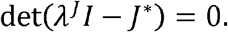

With the standard linear algebra technique for block matrix, when *N* < *M*, we can write the above equation as

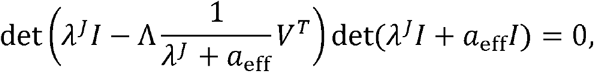

which can be further simplified as

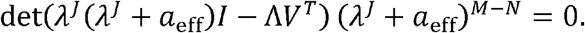

From this, we know there are *M* −*N* degenerate solutions *λ* ^*J*^ = − *a*_eff_, and another 2*N* solutions which can be obtained by solving

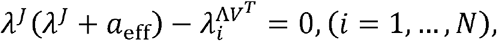

where 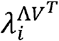 is the *i*th eigenvalue of Λ*V*^*T*^. For cases when *N* > *M*, similarly, we can break down the original determinant equation into

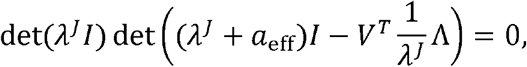

and then

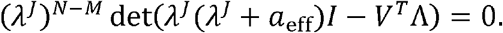

Besides the *N* − *M* degenerate solutions *λ* ^*J*^=0, we can have the rest 2*M* eigenvalues solved from

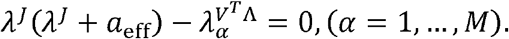

Similarly, we use 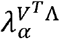 to denote the *α*th eigenvalue of Λ^*T*^*V*. To summarize, the positive eigenvalues of the Jacobian can be only obtained from the quadratic equations related to eigenvalues of Λ*V*^*T*^ or Λ^*T*^*V*.

Based on the scaling of different terms 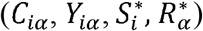, we know that the eigenvalues of Λ*V*^*T*^ or Λ^*T*^*V* have the magnetite 1/*M* or 1/*N*. This result is because ΛV^*T*^ has the same scaling behavior as *CC*^*T*^ (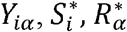 are of constant order), whose eigenvalues are proved to scale as 1/*M* ^54,68,69^ (since standard deviation of *C* element scales as 1/*M*). While the effective autoregulation is of constant order. In the limit of large *N* and large *M*, i.e., large complex system *a*_eff_ is much larger, and therefore, whether *λ* ^*J*^ can be positive is determined by whether the maximum (real part) 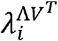 or 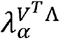 is positive. For large complex systems, we reduce the stability problem to a problem of studying Λ*V*^*T*^ or Λ^*T*^*V*.

Note that although we can reduce the study of Jacobian to that of Λ*V*^*T*^ or Λ*V*^*T*^ via assuming that that the reduction to Λ*V*^*T*^ or Λ^*T*^*V* is robustly correct when this assumption is not satisfied. A effective autoregulation matrix is proportional to an identity matrix, our numerical results show heuristic explanation is that the autoregulation terms can be all taken as infinity comparing to Λ and *V* in large communities, and thus are similar to being the same for Λ and *V*.

Next, we study the stability criterion for random matrix Λ*V*^*T*^ or Λ^*T*^*V*. If we have

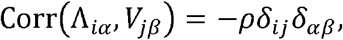

based on the random theory for non-symmetric Wishart matrix, the non-Hermitian Marchenko-Pastur law^54,68,69^ we can know Λ*V*^T^ has positive eigenvalues when *N* ≤ *M* once

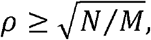

and *V*^*T*^ Λ has positive eigenvalues when *N* ≤ *M* if,

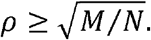

Therefore, we can combine the two cases to obtain a compact stability criterion as

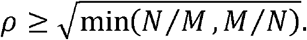

Note that the original (non-Hermitian) Marchenko-Pastur law requires the matrices to be i.i.d. and mean zero. Here, both Λ and *V* matrices have non-zero mean. However, the non-zero mean can be regarded as a low rank perturbation to the zero-mean part^83^, which would result in some negative outlier eigenvalues not affecting stability. Therefore, we can still apply the Marchenko-Pastur law here. Also, the presence of high-order regulation (*H*_*iαβ*_) will make elements within Λ correlated, not satisfying above assumption of correlation between elements exactly. But this correlation within Λ decays with increasing *M*, and thus the Marchenko-Pastur law prediction can be accurate in large communities.

We now try to express the abstract correlation coefficient by mechanistic quantities, e.g., the variation in yields, the statistics of production rates, as well as the statistics of high-order regulation effects. We estimate the correlation *ρ* for the equal concentration and equal abundance cases as previous works^41,84,85^, which is shown to work for general cases by numerical experiments. The standard deviation of Λ matrix element is approximated by

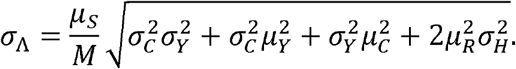

The standard deviation of *V* matrix element is approximated as

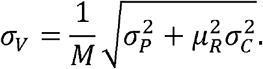

The covariance between Λ _*i α*_ and *V*_*iα*_ is

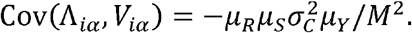

At the end, we have the estimated correlation as

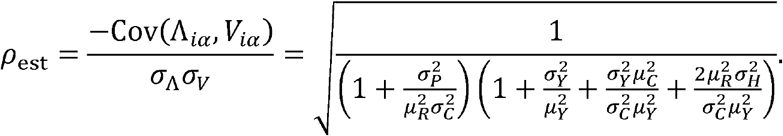

Substituting the estimated correlation into the stability criterion, we can obtain the two possible regions of diversity permitting stability,

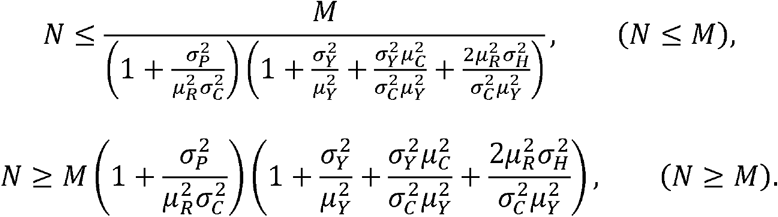

Heuristically comparing to May’s result (possible in the regime *N* ≤ *M*)^28^, where we have 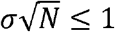 as the stability criterion, we can express the phenomenological interaction strength by the mechanistic variables as

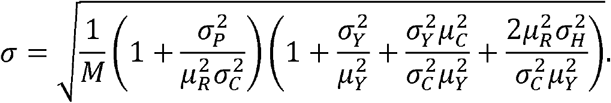

We use Julia to conduct numerical tests and run simulations and use Python for data analysis and drawing figures. Simulations are run on MIT Supercloud^86^. All codes are available at https://github.com/liuyz0/Critical-match. For a simulation process, we first determine the community diversities, i.e., the number of predators and that of prey. We then sample the parameters like consumption and growth rates based on given statistical parameters like mean and variance. We next sample the fixed point with given statistical parameters like mean and variance. Other parameters as mortalities and those in the prey/resource growth/supply function can be solved from the fixed point. We next can simulate the ordinary differential equations with various initial conditions. Time series outputted can give the information about survival diversity and fluctuating or not. After collecting all finial states from different initial conditions, if the final states are all stable, we can use principal component analysis to tell whether they are the same states (globally stable state) or not (alternative stable states). Except Fig. 1, which uses simulation data, we mainly did numerical experiments focusing on numerical analysis of the Jacobian without simulating the full dynamics. The sampling process is the same while stability can be directly read out from the eigenvalues of the numerically calculated local Jacobian eigenvalues.

