## Supplemental Information for "Ecosystem stability relies on diversity difference between trophic levels"

We studied the dynamics beyond the instability transition (Fig. 1b). We fixed predator diversity ($N=32$) and prey diversity ($M=64$), but changed standard deviation of yields $\sigma_{Y}$ from 0 to 1. Consumption rates $C_{i\alpha}$ are all sampled as i.i.d. and uniformly in $[0,1/M]$ (i.e., $\mathcal{U}(0,1/M)$). Predators and prey at the fixed point (i.e., $S^{*}$ and $R^{*}$) are sampled i.i.d. from $\mathcal{U}(0.01M/N,M/N)$ and $\mathcal{U}(0.01,1)$, respectively. We used logistic growth $h_{\alpha}\left( R_{\alpha} \right)=g_{\alpha}R_{\alpha}(K_{\alpha}-R_{\alpha})$, and sampling $g_{\alpha}$ i.i.d. from $\mathcal{U}(0.1,1)$. We sample yields as i.i.d. from Gaussian distribution with mean being $0.5$ and standard deviation $\sigma_{Y}$ (i.e., from $\mathcal{N}(0.5,\sigma_{Y}^{2})$). The rest of parameters can be solved at the fixed point. Given a community size, we sampled $100$ different communities with methods described, and for each community we randomly started from $50$ different initial conditions to see where the community will end up being. To best identify fluctuations while reducing simulation time cost, we check the convergence every 5e+3. We add a small constant dispersal 1e-7 to species during simulation. On the one hand, we can ensure positiveness of abundances. On the other hand, we can set a criterion for extinction, i.e., abundance lower than 1e-5 will be regarded as extinction, since the contribution of dispersal to abundance alone during each period can reach 1e-4. At the end of each 5e+3 period, we find the survival predators first, and check the average (over surviving species) relative difference between species abundances at the end and the mean values (over the last 1e+3 time). Manual inspection indicates the average relative difference for a converging community is typically smaller than 1e-3 while that for a fluctuating community is much larger, i.e., of the order 1e-1~1e+0. Therefore, we set the critical value as 1e-3 to distinguish stable and fluctuating communities. If a community is judged as being stable, we will end the simulation. Otherwise, we will continue a new 5e+3 period of simulation. The upper bound of the number of periods is 15. If a community does not converge after 15 periods, we will regard it as a fluctuating one. For fluctuation case, we can further use Julia package DynamicalSystems and the function lyapunov to identify the oscillatory attractor found is chaotic or not. With these procedures, we can obtain survival fraction, whether a system is fluctuating, and the type of fluctuation, given the parameters and one set of initial conditions. Since we want to distinguish alternative stable states and globally stable states, we need to test different initial conditions for a fixed set of parameters. Among all results obtained from different initial conditions, those go to stable states will be identified as globally stable as long as (i): there is no fluctuation starting from any initialization; and (ii): the stable states are the same fixed point. Otherwise, we will call these stable states as alternative stable states. We use principal component analysis (PCA) to tell whether different results in the space belongs to the same fixed point. The fractions of different types of attractors being observed are averaged over different communities and initial conditions.

We illustrated dynamical time series obtained from simulations (Fig. 1c). The top figure in Fig. 1c is obtained from a community with $20$ predators and $48$ prey. We set the variation of yields such that $\rho=0.8$. The consumption rates are sampled as in Fig. 1b. The fixed-point values of predators and prey are all $1$, and other parameters can be solved. The first straight part in the figure is not simulation but to show where the fixed point is. Then we set initial conditions of simulation or perturb the predators and prey out of the fixed point and start the simulation. The initial conditions for predators and prey are randomly sampled from $\mathcal{U}(0.5,1.5)$. The two middle figures in Fig. 1c are obtained from communities with $48$ predators and $48$ prey with the same sampling and simulation methods. The last figure is obtained from communities with $100$ predators and $48$ prey with the same sampling and simulation methods. In the last figure, to reduce confusions, we added a very small predator autoregulation $\epsilon_{i}=5\times{10}^{-4}$ to help the community to converge back to the original fixed point. If predator autoregulation is strictly zero when $N>M$, there will be a subspace rather than just a point (of predator and prey abundances) in which the community can fully coexist and stable (once the time trajectory hits the subspace, the community stay at the point hit). Each point in the subspace is marginally stable, with the eigenvectors with zero eigenvalues being basis of that subspace. We can regard this complex case as one more general stable and fully coexisting state. After adding very small predator autoregulation $\epsilon_{i}$, we found the communities always go back to the original fixed point with time ${\sim1/\epsilon}_{i}$ and the stability boundary does not change much.

We studied how predator diversity will affect stability (Fig. 1d). We fixed the number of prey ($M=48$) and the variation of yields (such that $\rho=0.8$), but changed predator diversity (from $N=1$ to $128$). We sample yields as i.i.d. from Gaussian distribution with mean being $0.5$ and standard deviation $\sigma_{Y}$ corresponding to $\rho=0.8$. The rest of parameters are sampled or solved the same as Fig. 1b. Simulation and data analysis are also the same as Fig. 1b.

In Fig. 2a, we sampled communities with different predator diversities ($N$ from $1$ to $64$) and prey diversities ($M$ from $1$ to $64$). Consumption rates $C_{i\alpha}$ are all sampled as i.i.d. and uniformly in $[0,1/M]$ (i.e., $\mathcal{U}(0,1/M)$). Predators and prey at the fixed point (i.e., $S^{*}$ and $R^{*}$) are sampled i.i.d. from $\mathcal{U}(0.01M/N,M/N)$ and $\mathcal{U}(0.01,1)$, respectively. We sample yields as i.i.d. from Gaussian distribution with mean being $0.5$ and standard deviation $\sigma_{Y}$ (i.e., from $\mathcal{N}(0.5,\sigma_{Y}^{2})$) and we fix yield variation such that correlation $\rho=0.8$. For each given community size ($N$ and $M$), we used the above sampling method to sample 20 communities, from which we can calculate the fraction of unstable ones. In Fig. 2b, we focused on $M=16, 32, 48$, and changed $N$ from $1$ to $64$ for more details: for each given system size, we then sampled 100 communities. In Fig. 2c, we used the same data generated in Fig. 2b, but plotted the figure with predator-prey ratio, $N/M$, rather than predator diversity, $N$.

In Fig. 3c, we varied yield variation and the predator diversity to generate the heatmap. The prey diversity is fixed as $32$, and the predator diversity change from $1$ to $128$ with a step size $1$. The consumption rates are sampled i.i.d. from $\mathcal{U}(0,1/M)$. The yields are sampled i.i.d. from $\mathcal{N}(0.5,\sigma_{Y}^{2})$, where $\sigma_{Y}$ is chosen such that $\rho_{\mathrm{est}}$ change from $0.001$ to $1$ with step size $0.001$. Other parameters are obtained the same as in Fig. 2. We then classified the communities based on $\rho_{\mathrm{est}}$ and $N/M$. In the figure, there are $100$ pixels along the $\rho_{\mathrm{est}}$ direction and $128$ pixels along the $N/M$ direction, such that each pixel contains $10$ communities with similar $\rho_{\mathrm{est}}$ and the same system size. We then calculate the heatmap value, fraction of unstable communities, for each pixel based on the $10$ communities in it. Fig. 3d is completely theoretical, plotting contour lines of $\rho_{\mathrm{est}}$ according to Eq. (6). The eigenvalues in Fig. 3e are obtained from matrices of the form $-AB^{T}/M$ (or $-B^{T}A/N$ when $N>M$), where $A$ and $B$ are $N\times M$ matrices with i.i.d. elements whose standard deviation is $1$. We show the examples when $M=64$. The correlation between $A_{i\alpha}$ and $B_{i\alpha}$ is fixed at $0.8$. Different colors refer to eigenvalues obtained from different sample matrices. Since the actual $\Lambda V^{T}$ differs from the standard $-AB^{T}/M$ by some rescaling that not changing the signs of eigenvalues (like rescaling the standard deviation to be $1$), it suffices to show the eigenvalues spectrum of $-AB^{T}/M$.

Fig. 4b is a panel in Fig. S6 (Fig. S6d), where the specific three-level Jacobian and numerical details will be explained. We fixed $N_{1}=32$, while change $M$ from 1 to 64 and $N_{2}$ from 1 to 64. Fig. 4b has $64\times64$ pixels, where value at each pixel is obtained by averaging over 10 communities. Fig. 4c used the same data from Fig. 4b but focused on cases having $M=1, 16, 32$.

We tested the validity of our theory for abiotic resources (Fig. S1). With the same setup as Fig. 3c, but changing the prey growth chemostat resource supply $h_{\alpha}\left( R_{\alpha} \right)=l_{\alpha}\left( \kappa_{\alpha}-R_{\alpha} \right)$, and sampling $l_{\alpha}$ i.i.d. from $\mathcal{U}(0.1,1)$, we can solve for the rest of parameters and calculate the Jacobian. The fraction of unstable communities is plotted, and the theoretical stability boundary (red line) is well supported by data. The fact that the stability criterion of large communities should not depend on resource supply/growth functions is implied by our derivation in Methods. To elaborate, the supply/growth functions affect only the specific forms of the effective autoregulation $a_{\mathrm{eff}}$. However, in the large community where $N$ and $M$ go to infinity, the effective autoregulation is sufficiently large regardless of specific forms and stability is only determined by $\Lambda V^{T}$, leading to the conclusion stability criterion will not be affected by the prey growth functions.

We studied how prey diversity will affect stability (Fig. S2). To that end, we fixed predator diversity $N=48$, and varied prey diversity $M$ from 1 to 128. The variation of yields is determined such that the correlation $\rho=0.8$. All other sampling and simulation methods are the same as Fig. 1d. We found there is also a re-entrant stability transition with respect to prey diversity that increasing $M$ first destabilizes but then stabilizes the communities.

We sought to discuss effects of other mechanisms on stability analysis first through a snapshot at a fixed community size. In Fig. S3a, we reviewed that variation of yields can lead to instability. We sampled communities with number of predators $N=16$, and number of prey $M=32$. Consumption rates $C_{i\alpha}$ are all sampled as i.i.d. and uniformly in $[0,1/M]$ (i.e., $\mathcal{U}(0,1/M)$). predators and prey at the fixed point (i.e., $S^{*}$ and $R^{*}$) are sampled i.i.d. from $\mathcal{U}(0.01M/N,M/N)$ and $\mathcal{U}(0.01,1)$, respectively. We sample yields as i.i.d. from Gaussian distribution with mean being $0.5$ and standard deviation $\sigma_{Y}$ (i.e., from $\mathcal{N}(0.5,\sigma_{Y}^{2})$) varying from $0$ to $1$ ($20$ values). The rest of parameters can be solved at the fixed point. For each given $\sigma_{Y}$, we sampled $100$ communities and record the stability via checking the Jacobian at the fixed point. Then, we can calculate the fraction of unstable communities at one $\sigma_{Y}$ value. In Fig. S3b and S3c, we used the same community size and sampling method as Fig. S3a. However, we set $\sigma_{Y}=0$ and varied $\sigma_{H}$ from 0 to 1 in Fig. S3b ($P$ were sampled from Gaussians) and $\sigma_{P}$ from 0 to 0.4 in Fig. S3c ($P$ were sampled from uniform distributions). We found stronger high-order dependence or cross-feeding (resource production) can lead to instability as predicted by theory (see Methods). After plotting fraction of unstable communities with respect to the correlation calculated, we found the instability transitions due to different mechanisms collapse, yet the one due to cross-feeding seems to be less consistent with the other two (Fig. S3d). We therefore conclude that the stability criterion based on correlation and diversity difference is robust to various mechanisms: different mechanisms alter the correlation in different manners while the stability criterion is not changed and works reasonably well.

We next discuss effects of other mechanisms on stability analysis systematically via results obtained by varying both community size and parameter statistics. We numerically studied the local Jacobian after adding different mechanisms and plot the fraction of unstable communities as a function of (estimated) correlation $\rho_{\mathrm{est}}$ and predator-prey ratio to compare with our stability criterion (Eq. (5)). Since we know the analytic dependence of how different mechanisms reduce the estimated correlation $\rho_{\mathrm{est}}$ (see Methods), when we sample a community with a desired $\rho_{\mathrm{est}}$value, we can balance the contributions from different mechanisms (in a way different mechanisms lead to similar decrease in $\rho_{\mathrm{est}}$). If we combine variation of yields, cross-feeding (or allowing resource production from consumers), and high-order prey regulations (i.e., the $H$ tensor), we found the theoretical stability boundary works robustly (Fig. S4a). However, if we compare the instability transition with those in communities only having yield variation (Fig. 4d), we found the transitions in Fig. S4a less sharp. We therefore conclude the robust validity of our theoretical stability boundary and the robustness of the re-entrant stability transition, but also found some mechanisms may make the theoretical prediction less accurate.

We next tried to study how the accuracy of the theoretical stability criterion is affected by different mechanisms. We tried to only include yield variation and high-order prey regulations, and found the stability transition is sharp near the theoretical boundary (Fig. S4b). If only yield variation and cross-feeding are considered, we obtained a fuzzy stability transition near the theoretical boundary (Fig. S4c). Combining all the evidence, we conclude that it is cross-feeding decreasing the accuracy of theoretical prediction.

We then discuss the robustness of the stability criterion theoretically. To obtain the stability criterion and the analytic form of $\rho_{\mathrm{est}}$, we ignored the distributions of fixed-point predators and prey, replacing the specific values by their mean. This operation can lead to accurate predictions if we only have yield variation. The reason is that the $\Lambda$ and $V$ matrices in this case can be rescaled by positive diagonal matrices to be independent of $S^{*}$ and $R^{*}$, which do not change stability. In other words, fixed point predator and prey abundances do not affect stability so we can assume fixed point values as long as they are positive. When we have high-order prey regulations $H$, the $\Lambda$ matrix cannot be rescaled to be independent of specific $R^{*}$ values. However, in large communities, only the mean of $R^{*}$ matters in determine the correlation between $\Lambda$ and $V$ (due to the summations $H_{i\alpha\beta}R_{\beta}^{*}$ and $H_{i\alpha\beta}R_{\alpha}^{*}$ and central limit theorem). We therefore still can get accurate predictions when having $H$ and using mean of $R^{*}$ to replace specific values in $R^{*}$. However, the introducing of cross-feeding (resource production) $P$ will make $V$ depend on $S^{*}$ in a way using mean of $S^{*}$ to replace specific values in $S^{*}$ affect the estimated correlation between $\Lambda$ and $V$. If we keep the true values of $S^{*}$ which are not identical, the $V$ matrix elements will not be i.i.d., and the non-Hermitian Marchenko-Pastur law may not directly apply. In fact, if we make $S^{*}$ values be uniformly identical, the stability boundary from simulation can be sharp which agrees with our theoretical boundary. In conclusion, resource production $P$ can violate the validity of approximation made during derivation or assumption needed for the random matrix theory used, and therefore leads to inaccuracy of our theoretical stability boundary.

We next discuss the effect of predator autoregulation ($\epsilon_{i}$ in Eq. (1)). By having the same setup as Fig. 4d but adding $\epsilon_{i}=0.05$ for all predators $i$, we observed that a lot of communities can be stabilized (Fig. S4d, where the dashed line is theoretical stability boundary without considering predator autoregulation). To understand the effects, we study the special example $\epsilon_{i}=\epsilon$ and the effective autoregulation for prey is also uniform as $a_{\mathrm{eff}}$. Following the same procedure in Methods, we can solve the eigenvalues of $J$, $\lambda^{J}$, from non-zero eigenvalues of $\Lambda V^{T}$ (or $V^{T}\Lambda$):

$$\left( \lambda^{J}+\epsilon\right)\left( \lambda^{J}+a_{\mathrm{eff}} \right)-\lambda_{i}^{\Lambda V^{T}}=0.$$

Given $a_{\mathrm{eff}}$ will be much larger than $\lambda_{i}^{\Lambda V^{T}}$ and note stability requires all $\lambda^{J}$ to have negative real parts, the approximate stability condition is that the maximum real eigenvalue of $\Lambda V^{T}$ should be smaller than $a_{\mathrm{eff}}\epsilon$. Since $a_{\mathrm{eff}}\epsilon>0$, the new critical correlation will be smaller than the original threshold $\min(\sqrt{N/M},\sqrt{M/N})$. The specific critical correlation will be related to the magnitude of $a_{\mathrm{eff}}\epsilon$ as well as that of eigenvalues of $\Lambda V^{T}$ which also depend on specific community size. It is then natural to see the stability transition in numerical results happens below the original boundary Eq. (5)). We therefore can reproduce the pattern for certain large correlation values that stability is irrelevant to diversities ($N$ or $M$), which is mentioned in the diversity-stability debate. We next ask whether there can still be re-entrant stability transition for small correlation values. Since the actual critical correlation value should be smaller than $\min(\sqrt{N/M},\sqrt{M/N})$, for a given correlation value can lead to instability, we can certainly change to sufficiently large $N/M$ or $M/N$ to make the communities stable. So, the re-entrant stability transition with respect to $N/M$ can still exit. In summary, we found predator autoregulation can stabilize the communities, making stability irrelevant to diversities under certain correlation values, but re-entrant stability transitions can exist for small correlation values.

We next discuss the finite size effects on stability transition. First, the random matrix result for $\Lambda V^{T}$ is rigorous for infinitely large communities, and therefore for finite size communities the instability transition will not be sharp near the predicted boundary. We copied Fig. 2c here (Fig. S5a), where the number of prey increases from $16$ (lightest dots) to $32$ (grey squares) to $48$ (dark triangles). Given the same ratio, $N/M$, communities with larger absolute size ($N$ or M) tend to have sharper transition near stability boundary predicted (dashed vertical lines). To further prove the concept, we did numerical tests to communities with $M=512$ (Fig. S5b), and found much sharper instability transitions. The second finite size effect is related to the reduction of the original Jacobian analysis to studying $\Lambda V^{T}$ (or $V^{T}\Lambda$). The eigenvalue spectrum of $\Lambda V^{T}$ is in an ellipse as predicted by the non-Hermitian Marchenko–Pastur law^54,68,69^ (Fig. S5c, where we used $N=32$, $M=128$, and correlation $\rho=0.8$). The analytic form of the ellipse in the complex plane is given by the equation right to Fig. S5c, where $x$ refers to the real axis and $y$ the imaginary axis. The analytic prediction (red curve in Fig. S5c) agrees well with numerical eigenvalue data (red dots in Fig. S5c). As is explained in Methods, we solve the eigenvalues of the original Jacobian $J^{*}$ via a quadratic equation from those of $\Lambda V^{T}$. To study this mapping from $\Lambda V^{T}$ eigenvalues to $J^{*}$ eigenvalues, we set different autoregulation values $a_{\mathrm{eff}}$ and solve $\lambda^{J}$ from $\lambda^{\Lambda V^{T}}$ (obtained in Fig. S5c) following formula in Methods. The solved $\lambda^{J}$ usually have four leaves on the complex plane (Fig. S5d, blue dots; the red curves are theoretical boundary for eigenvalues of $\Lambda V^{T}$). The real part of the center of $\lambda^{J}$ distribution is roughly ${-a}_{\mathrm{eff}}/2$. The values $\lambda^{\Lambda V^{T}}$ with real part greater than ${-a}_{\mathrm{eff}}^{2}/4$ will roughly be mapped to horizontal leaves of $\lambda^{J}$, and $\lambda^{\Lambda V^{T}}$ with real part smaller than ${-a}_{\mathrm{eff}}^{2}/4$ will roughly be mapped to vertical leaves of $\lambda^{J}$. If the effective prey autoregulation ($a_{\mathrm{eff}}$) is much greater than eigenvalues of $\Lambda V^{T}$, which is true in large communities due to our scaling of the mechanistic parameters with community size, the vertical leaves of $\lambda^{J}$ will vanish (Fig. S5e). In this case, the maximum $\lambda^{J}$ is the one on the real axis and is determined by the maximum real eigenvalue of $\Lambda V^{T}$, yielding our main results reported. However, when the community size is not large, $a_{\mathrm{eff}}$ will not be guaranteed to be much greater than eigenvalues of $\Lambda V^{T}$. If $a_{\mathrm{eff}}$ is small comparing to the eigenvalues of $\Lambda V^{T}$, the horizontal leaves of $\lambda^{J}$ will shrink or vanish and the eigenvalues with maximum real part come from vertical leaves (Fig. S5f). The stability now depends on how wide the vertical leaves are along the real axis (related to how wide $\Lambda V^{T}$ eigenvalues are along the imaginary axis) and the center of the vertical leaves on the real axis (related to $a_{\mathrm{eff}}$). If Jacobian eigenvalues from the vertical leaves become unstable first, the whole community will be unstable while the reduced inter-predator interaction $\Lambda V^{T}$ is still stable. We should interpret this new pattern of losing stability as a failure of autoregulation to sustain predator-prey interactions. A simple example of this instability is Lotka or Volterra's original prey-predator model^78,79^, where there is only one prey species and one predator species, and the inter-predator interaction is always stable $\Lambda V^{T}$ is a negative scalar number) but the community can be unstable (in neutrally stable limit cycles). We therefore discussed possible inaccuracies in our theory regarding small-sized communities. The first concerns the inaccuracy of infinite size random matrix theory, which yields a less sharp instability transition in small communities than predicted. The second involves potential small effective autoregulation, leading to another way of losing stability, even though reduced inter-predator (or inter-prey) interactions remain stable.

We next discussed stability in communities with more than two levels and the generality of our conclusions drawn from communities with only two levels. Our preliminary exploration focused on three-level communities, a bottom level with $M$ prey species (level $0$), a middle level with $N_{1}$ predator species (level $1$), and a top level with $N_{2}$ apex predator species (level $2$). And we assume the simple case where level $\mathcal{l}$ only interact with level $\mathcal{l\pm}1$. Comparing to the two-level community dynamics, the only difference is we added a new level grow on predators. Similar to equations in Methods, the Jacobian should have the form as Fig. S6a. Here, $V^{\mathcal{(l)}T}$ encodes how level $\mathcal{l+}1$ affects level $\mathcal{l}$, $\Lambda^{\mathcal{(l)}}$ encodes how level $\mathcal{l-}1$ impacts level $\mathcal{l}$, and $a^{\mathcal{(l)}}$ denotes effective autoregulation of level $\mathcal{l}$. For simplicity, we assume uniform autoregulation within each level to observe some patterns qualitatively (note that in the conventional consumer-resource models, $a^{\mathcal{(l)}}=0$ for $\mathcal{l>}0$). For the numerical results shown in the following, we fixed $N_{1}=32$ and varied $N_{2}$ and $M$. We sampled the matrix elements for both $\Lambda^{\mathcal{(l)}}$ and $V^{\mathcal{(l)}}$ from Gaussian $\mathcal{N}(3,1)$. The mean value does not matter as long as it is positive, which contribute to outlier eigenvalues always stable. The variance can matter when autoregulation is too small. We also fixed the correlation between $\Lambda^{\mathcal{(l)}}$ and ${-V}^{\mathcal{(l)}}$ as $0.8$ in the tests and only studied how various community sizes change community stability. If we have autoregulation in all levels ($a^{\mathcal{(l)}}=1$ for all $\mathcal{l}$), we find the communities are all stable (Fig. S6b). This is consistent with the finding in two-level communities where if autoregulation exists for all levels, there can be correlation values for which the communities are always stable (Fig. S4d). If only level $2$ does not have autoregulation ($a^{\mathcal{(l)}}=1$ for $\mathcal{l=}0,1$ and $a^{(2)}=0$), we found stability is determined by diversity difference between the higher two levels, i.e., the ratio $N_{2}/N_{1}$ (Fig. S6c). We can see the stability problem for these three-level communities reduce to the stability problem for the higher two levels. If only level $1$ does not have strong autoregulation ($a^{\mathcal{(l)}}=1$ for $\mathcal{l=}0,2$ and $a^{(1)}=0$), we found stability is determined by ($M+N_{2})/N_{1}$: the community is stable if total diversity of level $0$ and $2$ is very different from diversity of level $1$ (Fig. S6d, which is also Fig. 4b). If only level $0$ does not have autoregulation ($a^{\mathcal{(l)}}=1$ for $\mathcal{l=}1,2$ and $a^{(0)}=0$), the stability will be determined by $M/N_{1}$, i.e., it reduces to the stability problem of the lower two levels (Fig. S6e). If only level $2$ has autoregulation ($a^{\mathcal{(l)}}=0$ for $\mathcal{l=}0,1$ and $a^{(2)}=1$), the communities are almost always unstable as the first two levels may always be unstable (Fig. S6f). If only level $1$ has autoregulation ($a^{\mathcal{(l)}}=0$ for $\mathcal{l=}0,2$ and $a^{(1)}=1$), the stability will be determined by ($M+N_{2})/N_{1}$ suggesting we may imagine level $0$ and $2$ as one level consider a two level stability problem between this joint level and level $1$ (Fig. S6g). If only level $0$ has autoregulation ($a^{\mathcal{(l)}}=0$ for $\mathcal{l=}1,2$ and $a^{(0)}=1$), the communities are almost all unstable, and stability only shows up when $N_{2}/N_{1}$ is small while $M$ sufficiently differs form $N_{1}$ (Fig. S6h). If all three levels do not have autoregulation ($a^{\mathcal{(l)}}=0$ for all $\mathcal{l}$), the communities will be always unstable unless $M$ and $N_{2}$ are very small, which may be due to finite size effect (Fig. S6i). Combining all the patterns, we found that we need at least one level to have weak enough autoregulation to have instability in large communities. And difference between diversity of levels having strong autoregulation and that of other levels seems to control the re-entrant stability transition. In general, the idea stability depends on diversity difference among levels rather than total diversity of the community still hold while the specific quantification of diversity difference as well as the threshold value may be different for more complex situations.

We need to note that cases studied here are preliminary. To have non-trivial instability transitions in communities with more than two levels, the autoregulation for different levels of predators is necessary whose form or scaling need to be justified. Also, the correlation between $\Lambda^{\mathcal{(l)}}$ and ${-V}^{\mathcal{(l)}}$ for different $\mathcal{l}$ can be different. There can also be more interactions between level $\mathcal{l}$ and levels beyond $\mathcal{l\pm}1$. In general, the stability problem for communities with more than two levels can be much more complex in terms of setup and analysis, which need future justifications and explorations.

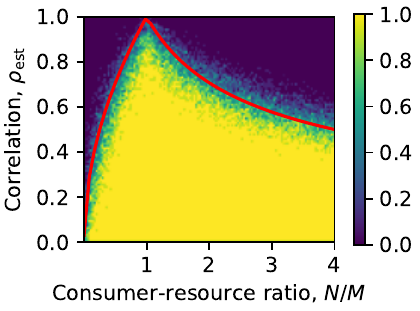

Fig. S1 | The stability criterion is valid for chemostat resource supply (consumer here is equivalent to predator, and resources equivalent to prey). The heatmap value refers to fraction of unstable communities at the given position and is obtained from numerically sampled communities with $M=32$ and $N$ varying from $1$ to $128$. There are $128\times100$ pixels, and each pixel has $10$ communities fell in, from which the unstable community fraction can be calculated.

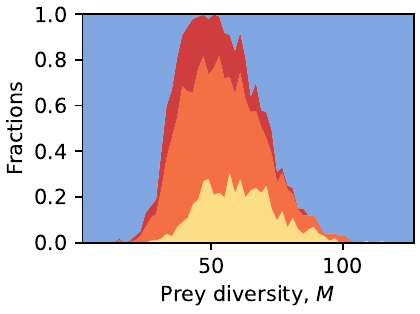

Fig. S2 | There is also a re-entrant stability transition with respect to prey diversity. Only varying prey diversity, $M$, we can observe a similar re-entrant stability transition, i.e., increasing prey diversity first destabilizes but then stabilizes communities. The color encoding for different dynamical behaviors is the same as Fig. 1, and we changed $M$ from $1$ to $128$ while kept $N=48$.

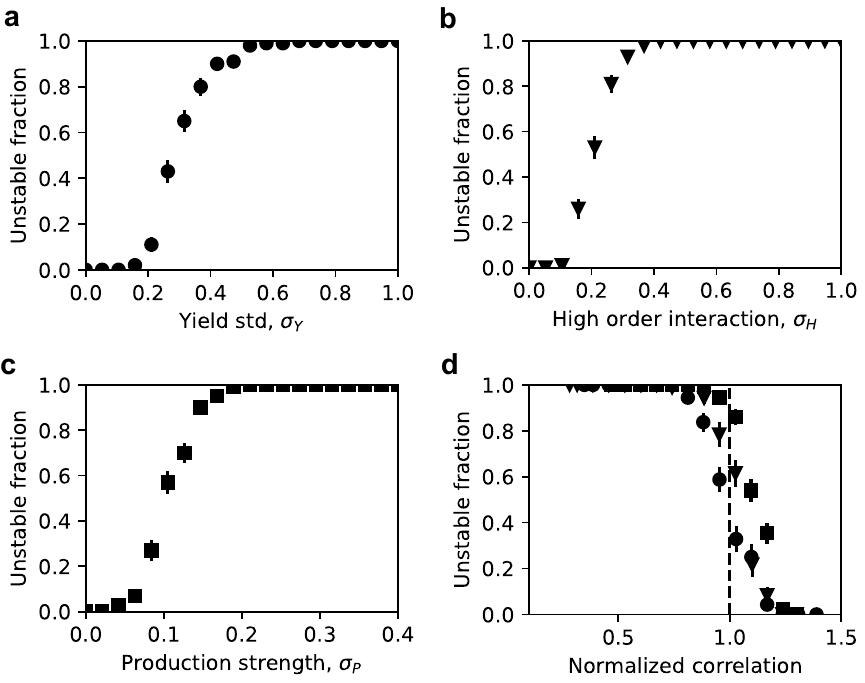

Fig. S3 | Our stability criterion unifies instability transitions due to different mechanisms when community sizes are fixed. (a) Only varying standard deviation of yields, $\sigma_{Y}$, we see the communities lose stability with increasing $\sigma_{Y}$ (communities have $N=16$, $M=32$). (b) Only varying variance of high order interactions, we observe one instability transition ($N=16$, $M=32$). (c) Only increasing strength of resource production (cross-feeding), we can observe one instability transition ($N=16$, $M=32$). (d) After plotting the data points in (a)-(c) with respect to normalized correlation $\sqrt{2}\rho_{\mathrm{est}}$ (see Methods for formula of $\rho_{\mathrm{est}}$), the instability transitions due to different mechanisms collapse. Note the theoretical stability criterion is $\rho_{\mathrm{est}}>\sqrt{N/M}=1/\sqrt{2}$ since $N=16$ and $M=32$, the instability transition is predicted to happened near $\sqrt{2}\rho_{\mathrm{est}}=1$ (dashed line). Therefore, the instabilities due to different mechanisms can be interpreted to reduce alignment between how predators affect prey and how prey affect predators back, making predators more likely to encroach upon niches (growth promoting prey) of other predators.

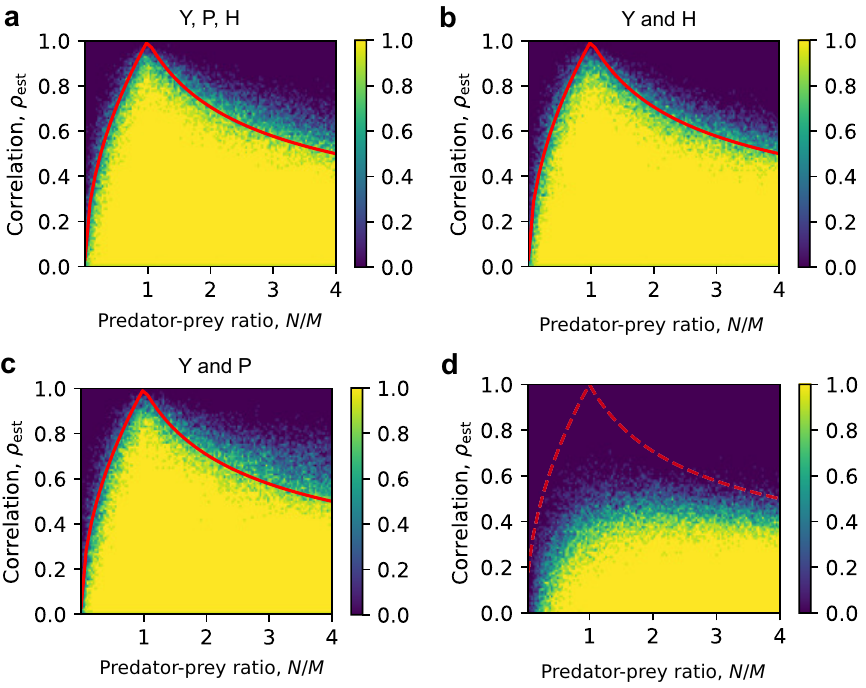

Fig. S4 | Re-entrant stability transition is robust to various mechanisms. (a) The stability criterion Eq. (5) depicted by the red curve well identifies instability transition after allowing yield variation, cross-feeding (or resource production by consumers), and high-order dependence of predator growth on prey. The heatmap value refers to the fraction of unstable communities. (b) The stability criterion Eq. (5) is accurate in identifying the instability transition if we only allow yield variation and high-order effects. (c) However, allowing cross-feeding will make the stability criterion Eq. (5) less accurate since the existence of cross-feeding can violate the validity of approximation made during derivation (assumption needed for the random matrix theory used). The re-entrant stability transitions still exist. (d) Predator autoregulation can stabilize the communities reducing the critical correlation value from $\min\left\{ \sqrt{N/M},\sqrt{M/N} \right\}$ (dashed red curve). Therefore, stability can be irrelevant to diversities ($N, M$) under certain large correlation values. But re-entrant stability transitions can robustly exist for smaller correlation values.

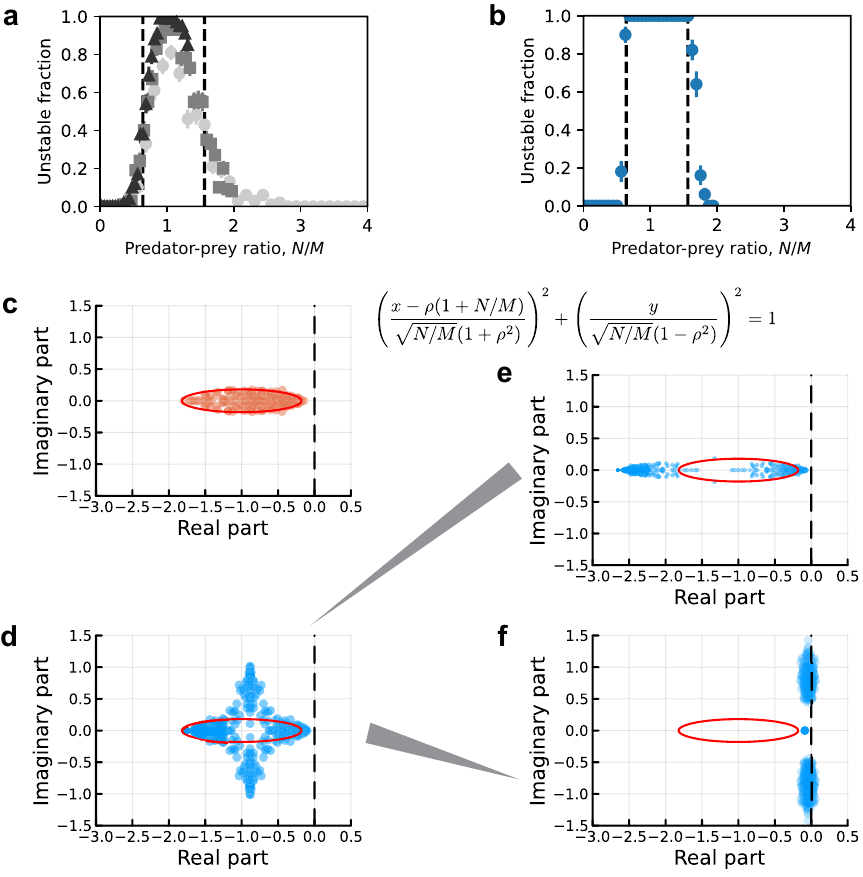

Fig. S5 | Stability of finite (or small) size communities may not be accurately predicted by stability of inter-predator interactions. (a) Increasing community size will make the prediction based on large random matrix more accurate (instability transition will be sharper). The number of prey increases from 16 (lightest dots) to 32 (grey squares) to 48 (dark triangles). (b) For communities with 512 prey, i.e., very large community sizes, the instability transition is sharp and close to be discontinuous. (c) The eigenvalue spectrum (red points) of $\Lambda V^{T}$ is in an ellipse (the red curve; the equation of this ellipse is given right to (c)). Data are generated when $N=32$, $M=128$, and correlation $\rho=0.8$. (d) The eigenvalues of Jacobian (blue dots) have four leaves and can be solved from those of $\Lambda V^{T}$ (red curve is the theoretical boundary of the eigenvalues of $\Lambda V^{T}$). (e) When effective autoregulation is large comparing to the eigenvalues of $\Lambda V^{T}$, the vertical leaves of Jacobian eigenvalues vanish and stability is determined by eigenvalues in the horizontal leaves, where the maximum one is determined by the maximum real eigenvalue of $\Lambda V^{T}$. (f) When autoregulation is small, the vertical leaves of Jacobian eigenvalues can lose stability first, which is a new way of losing stability while the reduced inter-predator interaction $\Lambda V^{T}$ is stable.

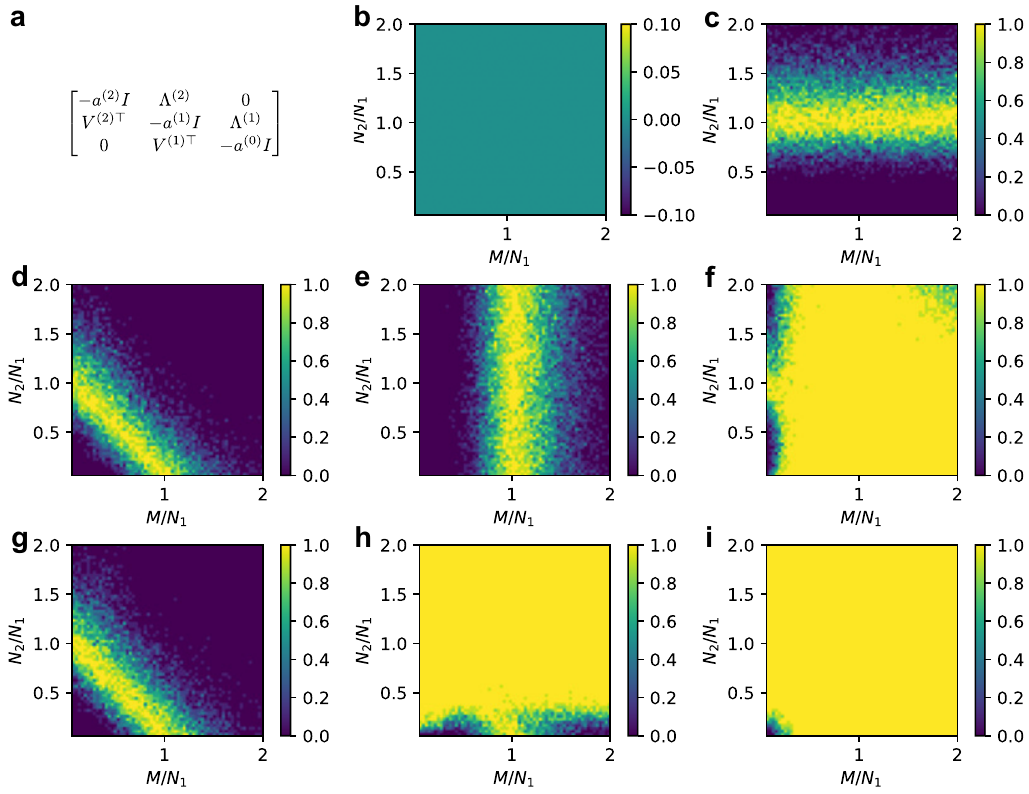

Fig. S6 | Stability of special three-level communities depends on diversity differences among levels. (a) The Jacobian of three-level communities: a level with $M$ prey (level $0$), a level with $N_{1}$ predator species (level $1$), and a level with $N_{2}$ apex predator species (level $2$). Matrix $V^{\mathcal{(l)}T}$ encodes how level $\mathcal{l+}1$ affects level $\mathcal{l}$, $\Lambda^{\mathcal{(l)}}$ encodes how level $\mathcal{l-}1$ impacts level $\mathcal{l}$, and $a^{\mathcal{(l)}}$ denotes effective autoregulation of level $\mathcal{l}$. (b) If we have non-negligible autoregulation in all levels ($a^{\mathcal{(l)}}=1$ for all $\mathcal{l}$), we find the communities are all stable. Heatmap value here and after refers to the fraction of unstable communities. (c) If only level $2$ does not have strong autoregulation ($a^{\mathcal{(l)}}=1$ for $\mathcal{l=}0,1$ and $a^{(2)}=0$), we found stability is determined by diversity difference between the higher two levels. (d) If only level $1$ does not have strong autoregulation ($a^{\mathcal{(l)}}=1$ for $\mathcal{l=}0,2$ and $a^{(1)}=0$), we found stability is determined by $\left( M+N_{2} \right)/{N_{1}}$. (e) If only level $0$ does not have strong autoregulation ($a^{\mathcal{(l)}}=1$ for $\mathcal{l=}1,2$ and $a^{(0)}=0$), the stability will be determined by $M/{N_{1}}$. (f) If only level $2$ has non-negligible autoregulation ($a^{\mathcal{(l)}}=0$ for $\mathcal{l=}0,1$ and $a^{(2)}=1$), the communities are almost always unstable. (g) If only level $1$ has autoregulation ($a^{\mathcal{(l)}}=0$ for $\mathcal{l=}0,2$ and $a^{(1)}=1$), the stability will be determined by $\left( M+N_{2} \right)/{N_{1}}$. (h) If only level $0$ has non-negligible autoregulation ($a^{\mathcal{(l)}}=0$ for $\mathcal{l=}1,2$ and $a^{(0)}=1$), the communities are almost all unstable. (i) If all three levels do not have sufficient autoregulation ($a^{\mathcal{(l)}}=0$ for all $\mathcal{l}$), the communities will be always unstable.
